# Split-Transformer Impute (STI): A Transformer Framework for Genotype Imputation

**DOI:** 10.1101/2023.03.05.531190

**Authors:** Mohammad Erfan Mowlaei, Chong Li, Oveis Jamialahmadi, Raquel Dias, Junjie Chen, Benyamin Jamialahmadi, Timothy Richard Rebbeck, Vincenzo Carnevale, Sudhir Kumar, Xinghua Shi

**Affiliations:** Computer & Information Sciences, Temple University, 1925 N. 12th Street, Philadelphia, 19122, PA, USA; Department of Molecular and Clinical Medicine, Institute of Medicine, Sahlgrenska Academy, Wallenberg Laboratory, University of Gothenburg, Gothenburg, Sweden; Department of Microbiology and Cell Science, University of Florida, 1355 Museum Dr, Gainesville, 32603, FL, USA; Computer Science and Technology, Harbin Institute of Technology, Shenzhen University Town, Shenzhen, 518055, Guangdong, China; David R. Cheriton School of Computer Science, University of Waterloo, 200 University Avenue West, Waterloo, N2L 3G1, ON, CA; Division of Population Sciences, Dana-Farber Cancer Institute, 450 Brookline Ave, Boston, 02215, MA, USA; Department of Epidemiology, Harvard T. H. Chan School of Public Health, 677 Huntington Ave, Boston, 02115, MA, USA; Institute for Genomics and Evolutionary Medicine, Temple University, 1925 N. 12th Street, Philadelphia, 19122, PA, USA; Institute for Computational Molecular Science, Temple University, 1925 N. 12th Street, Philadelphia, 19122, PA, USA; Department of Biology, Temple University, 1925 N. 12th Street, Philadelphia, 19122, PA, USA

**Keywords:** Genotype, Structural variation, Imputation, Deep learning, Transformer

## Abstract

**Motivation:** Despite recent advances in sequencing technologies, genome-scale datasets continue to have missing bases and genomic segments. Such incomplete datasets can undermine downstream analyses, such as disease risk prediction and association studies. Consequently, the imputation of missing information is a common pre-processing step for which many methodologies have been developed. However, the imputation of genotypes of certain genomic regions and variants, including large structural variants, remains a challenging problem.

**Results:** Here, we present a transformer-based deep learning framework, called a split-transformer impute (STI) model, for accurate genome-scale genotype imputation. Empowered by the attention-based transformer model, STI can be trained for any collection of genomes automatically using self-supervision. STI handles multi-allelic genotypes naturally, unlike other models that need special treatments. STI models automatically learned genome-wide patterns of linkage disequilibrium (LD), evidenced by much higher imputation accuracy in high LD regions. Also, STI models trained through sporadic masking for self-supervision performed well in imputing systematically missing information. Our imputation results on the human 1000 Genomes Project show that STI can achieve high imputation accuracy, comparable to the state-of-the-art genotype imputation methods, with the additional capability to impute multi-allelic structural variants and other types of genetic variants. Moreover, STI showed excellent performance without needing any special presuppositions about the patterns in the underlying data when applied to a collection of yeast genomes, pointing to easy adaptability and application of STI to impute missing genotypes in any species.

## 1 Introduction

Genetic and genomic studies, such as linkage analysis, genome-wide association study (GWAS), and polygenic risk score (PRS) estimation, enable us to dissect the genetic architecture of complex traits and diseases [1]. In recent years, whole-genome sequencing (WGS) platforms and techniques have substantially improved and become increasingly cost-effective, resulting in the accumulation of large collections of genotypes and deeper insights into the genetic architecture of various traits and diseases.

Although the resolution of genotyping has steadily improved over time, genotype data still contain many missing values and untyped loci [2]. The missing data may decrease statistical power in disease association studies and causal variant discovery [3–5]. Causes of missing genotypes include the difficulty in sequencing rare alleles [6–8], failure of experimental assays, genotype calling errors, and differences in densities and properties of genotyping platforms [3]. As such, genotype missingness, as depicted in Figure 1, can be classified into two distinct categories: sporadic missingness, where for each site/segment, some values could be absent, and systematic missingness, in which some genomic loci or segments are not genotyped. These challenges in handling missing data are further compounded when considering different types of genetic variation in addition to Single Nucleotide Variants (SNVs). Compared with SNVs, Structural Variations (SVs) pose greater challenges in genotype calling and imputation due to their increased complexity, limitations of current sequencing technologies, extensive allelic diversity, and their variable frequencies within populations [9, 10]. Moreover, SVs can have a more significant impact on genetic diseases than SNVs, so their accurate imputation can lead to enhancements in disease association studies [11].

**Fig. 1.**
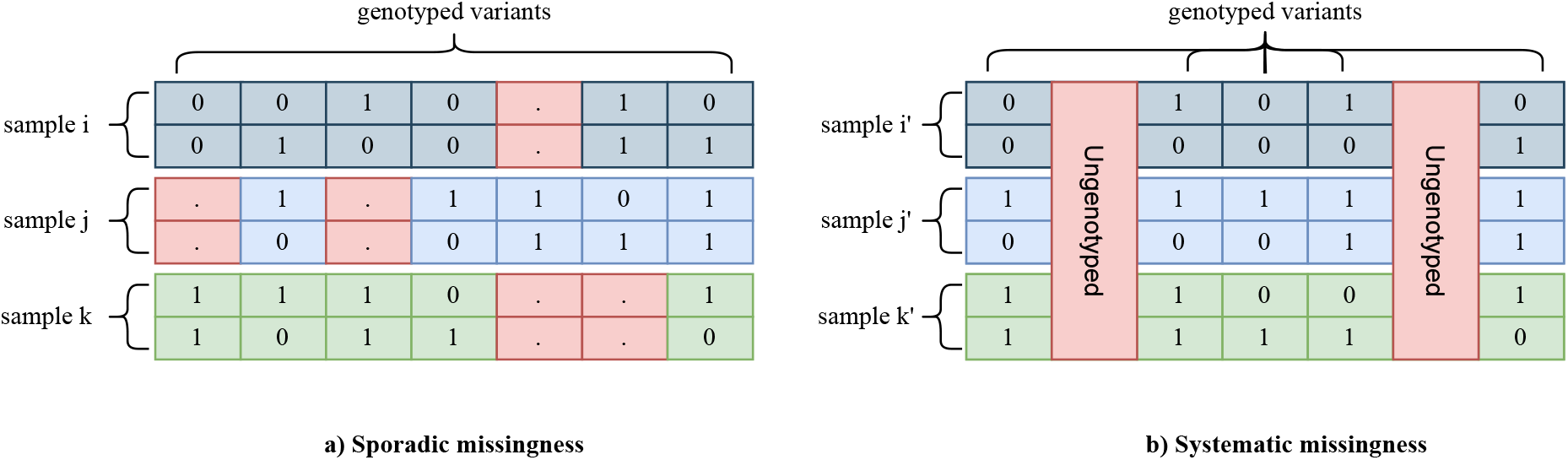
Missing genotype classification. *a. Sporadic missingness:* This category is commonly associated with methods in genotype calling and assay failures. *b. Systematic missingness:* Differences in sequencing resolution are common causes of this type of missingness.

Consequently, there is a common need for reliable imputation of genotypes using computational methods. Imputation is the process of inferring missing values in the data based on the information already present in the dataset, such as the density and distribution of bases and structural variants within and among sequences in the dataset. Imputation of missing data in genomics needs specialized methods because genomic information is inherently different from data in many other domains, such as vision or natural language processing. Curse-of-dimensionality, linear and non-linear correlations among the variants [12], and shared segments of sequences due to common descent are among the unique characteristics of genotype data. Furthermore, given the multifaceted nature of genotype imputation, it is well-recognized that no single method can serve as a universal solution. Consequently, it is a common practice in the field to utilize multiple imputation tools for a particular study.

Widely-used imputation methods often require a reference panel of genome sequences to impute missing information in sequences, assuming that the missing information comes from the same ancestry patterns as those in the reference panel [13]. These methods utilize Hidden Markov Models (HMMs), graphical models, and haplotype-cluster algorithms to impute missing values [14]. For example, Minimac4 [15], the most recent version of MACH [16], uses an HMM. For each individual, Minimac4 updates the phase iteratively in both directions based on haplotypes in the reference panel and neighboring loci in the individual. It splits sequences into overlapping chunks in order to reduce memory consumption and make the model scalable. Similarly, Shapeit5 [17], IMPUTE2 [18] and BEAGLE [19] also employ HMMs to perform imputation. The Haplotype-clustering algorithm is utilized in the fastPHASE [20] in order to cluster haplotypes in an SNV-wise manner and impute missing values per locus. GLIMPSE2 [21] uses a HMM in order to genotype low-coverage whole-genome sequencing (WGS) data.

Deep Learning (DL) methods have been recently introduced for genomic imputations. Sparse Convolutional Denoising autoencoder (SCDA) is used in [14] to impute missing data in the Human Leukocyte Antigen (HLA) region on chromosome 6 and yeast [22] genotypes. In [3] an improvement in SCDA training is proposed to improve the performance. Similarly, [23] used autoencoders on identified linkage disequilibrium (LD) blocks as well as focal loss to improve the performance. RNN-IMP [24] utilizes recurrent neural networks (RNNs) and augments the samples using recombination and mutation in order to impute systematic missingness in genotype data. GRUD [25] utilizes RNNs in an adversarial training schema. DEEP*HLA [26] uses a convolutional neural network to perform imputation on pre-phased genotypes at the gene level. Inspired by DEEP*HLA, HLARIMNT [27] uses transformers for the same task.

Though the overall performance of existing imputation methods for genomic data is generally good, most of them cannot directly handle multi-allelic variants. Also, their performances have not been evaluated for imputing SVs to the best of our knowledge. Also, the training of DL models [3, 14, 23, 26] is generally slow, and they need many more samples to perform as well as methods based on HMMs, such as Minimac4 [15] and Beagle5.4 [19]. Although RNN-IMP and GRUD [24, 25] addressed this performance disparity, these methods require retraining when the variant sets in the target are different from those in the training set. They are also not designed to handle sporadic missingness (Figure 1.a). Finally, the rest of the existing DL methods rely on convolutional neural networks (CNNs), which excel at exploiting local patterns but do not exhibit a robust mechanism to effectively capture pairwise correlations among local and distant markers simultaneously, such as the presence of LD blocks in genotypes.

An effective solution to this is the attention mechanism in the transformer architecture, capable of capturing local and distal interactions in genomes[28]. The attention mechanism in DL mimics visual attention to focus on specific parts of pictures [29, 30] by calculating importance scores among genomic loci. Therefore, attention can capture global interactions amongst markers. Transformers utilize multihead attention to capture intricate and multi-level interactions among the variants. AlphaFold2 [31] and ESMFold [32] are successful examples of transformers in biological sequence analysis.

In this article, we present a novel genotype imputation model, STI, based on the attention mechanisms in a transformer framework. Our model utilizes attention to capture patterns among SNVs and SVs in the genome collections analyzed. We found STI to achieve high imputation accuracy at a modest memory consumption cost, achieved by dividing the data into chunks (following [15]) that enables efficient application of STI to long sequences. Furthermore, STI needs to be trained only once, unlike other DL models, following that the imputation times in STI are faster than classical methods (Table 11 in the Supplement). In brief, our study makes the following key contributions.

- We propose a DL transformer framework termed STI, designed to specifically address the genotype imputation problem.
- STI imputation does not need a standard reference panel, which makes it more generally applicable to various data formats.
- STI excels at SV imputation, where the variants harbor a higher degree of complexity while achieving comparable performance to competing imputation models for SNV imputation.
- We analyze the effect of different masking rates on building better imputation models and explain the reasons for the STI improvements.

## 2 Results

### 2.1 Overview of the study

In this section, first, we present the results of our empirical study to find the optimal masking percentage (masking rate, MaskR) for STI training in order to eliminate the need for building imputation models for specific missing rate (MissR) in the target dataset, which is often required by some other machine learning approaches [3, 14]. After that, we present results for sporadic missingness imputation on the yeast dataset, SVs in human chromosome 22, and extensive SVs dataset (see Subsection 4.1 in Methods for the dataset details). We benchmarked STI’s performance against classical imputation methods (Beagle5.4 and Minimac 4.1.4), deep learning models (SCDA, AE, DEEP*HLA), and a variant of STI that uses no embedding (STI-NE). The details of STI architecture and aforementioned methods are discussed in Subsections 4.3 and 4.4, respectively.

### 2.2 Optimal masking rate analysis

For this analysis, we used the HLA region on chromosome 6 from the human 1000 Genomes Project and performed a 3-fold cross-validation on the data. The aim was to examine the relationship of MaskR for the training set with varying MissR in target sets. The results are presented in Figure 2 in which the results on the left/right column belong to the validation/test set, respectively.

**Fig. 2.**
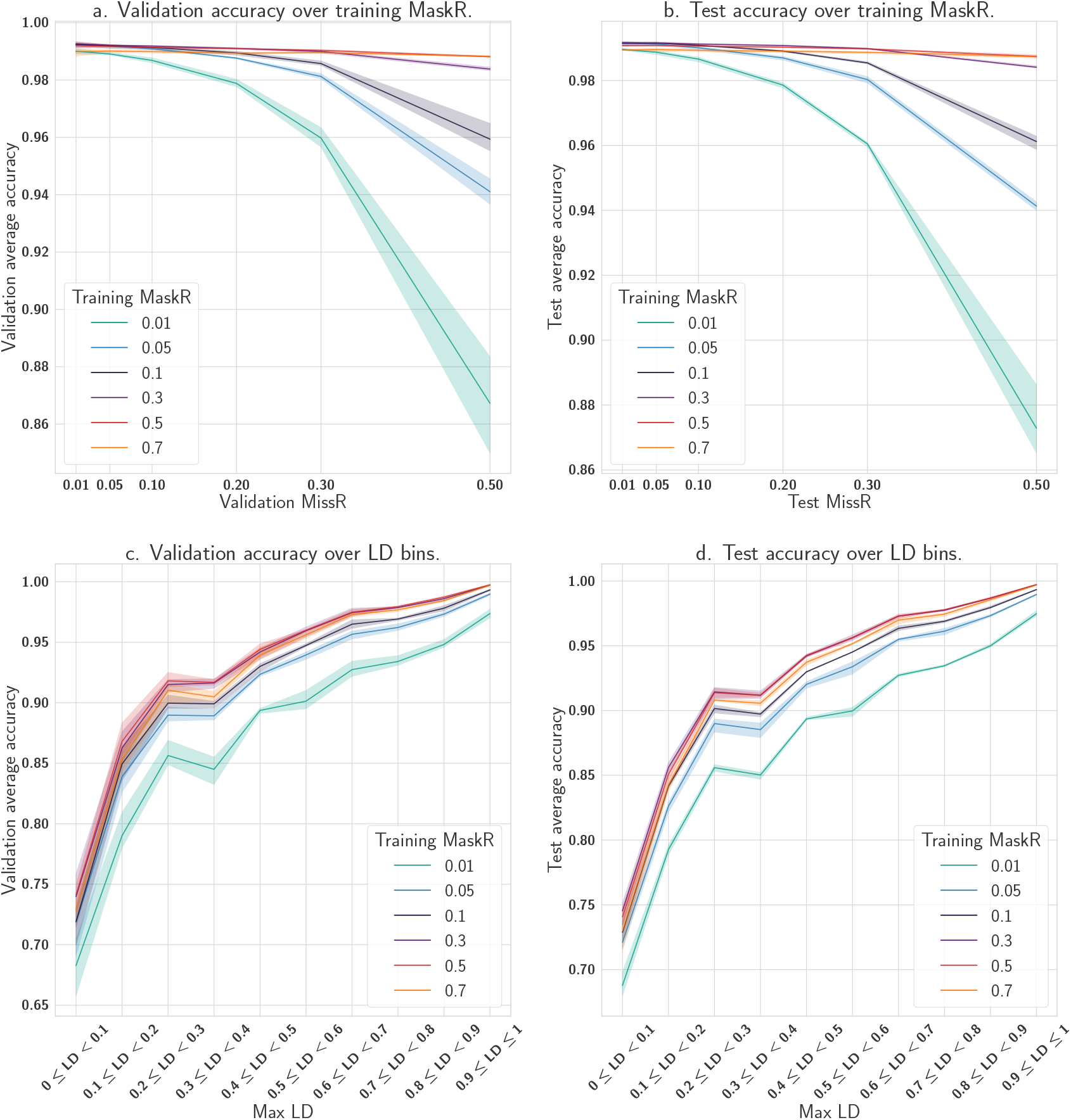
Average accuracy over 3-fold cross-validation for validation and test sets in the HLA dataset using different MaskR values during training. *a*. and *b*. A breakdown of average accuracy for various MissRs of validation/test set when the model is trained using different MaskR values. The patterns show that a model trained using a higher MaskR is more robust across different target MissRs. *c*. and *d*. Average accuracy for validation/test sets over 3 folds and MissRs of 0.01, 0.05, 0.1, 0.2, 0.3, and 0.5 calculated for various LD bins. The trend suggests that a higher MaskR increases the performance across LD bins, which could be attributed to the impact of MaskR on STI to learn LD patterns comprehensively. When MaskR is low, STI imputations do not benefit from the LD patterns present and, thus, STI does not learn the majority of pairwise correlations (LD) among the variants. Consequently, STI is not able to infer the missing value using all possible information in the respective LD block of the target variant.

Figures 2 *a & b* show that the performances of STI models trained using MaskR of 0.5 and 0.7 were high for imputations in which MissRs were up to 0.5. Therefore, a single STI model, trained with a MaskR of 0.5, could be used for a variety of research datasets as long as the MissR is less than 0.5. However, STI models trained with MaskR *>* 0.5 is needed for reliable imputations when the target datasets have more than 50 percent missing variants. Therefore, we recommend STI models trained with a MaskR of 0.5 for imputing sporadically missing variants and a higher MaskR (up to 0.8, as indicated by our empirical studies data) for other datasets.

Figures 2 *a & b* also show that when the target MissR is sufficiently low, the performance gap of the imputation models is not discernible. The performance gap becomes evident with a MissR of 0.2 or higher.

The underlying cause of this observation is that when the MissR is extremely low, a sufficient number of variants in LD with the target variant are readily available, making predictions less challenging for all the models. Conversely, a large MissRs means that the amount of information from LD blocks diminishes, presenting a greater challenge to the imputation model.

Figures 2 *c & d* show that, generally, STI models trained with lower MaskR will produce poor performance for imputing missing SNVs located in regions with high LD. For instance, variants in regions with LD = 0.01 have the lowest accuracy for all the masking percentages. Additionally, these results indicate that it is easier to predict missing data in high LD regions compared to low LD regions, which aligns well with biological expectations that low LD regions do not benefit from additional information (LD) available for better imputation of high LD regions. These trends suggest that the use of a low MaskR prevents the model from learning LD patterns, resulting in a worse performance. In other words, the model training needs to effectively disturb the LD blocks (and other latent patterns among variants) to capture direct and indirect correlations and haplotypes. Consequently, MaskR of 0.5 and higher provides robust results across a large range of target MissR values.

### 2.3 The relative performance of STI for sporadic missingness

For each dataset in this experiment, we performed a 3-fold cross-validation where missing values were introduced using fixed random seeds to ensure reproducibility of results across experiments and methods. The missing values were distributed randomly according to one of three strategies: uniformly, based on Minor Allele Frequency (MAF), or based on LD. These methods were chosen to ensure that missing values are representative of the data distribution in different biological aspects. Further details on these procedures can be found in the methods section. In all of the experiments, missing positions in the test sets were the same for all the methods.

The overall results for the yeast and chromosome 22 datasets are presented in Table 1. The numerical values in this table indicate the average of the metric values on the test sets in a 3-fold cross-validation.

**Table 1.**
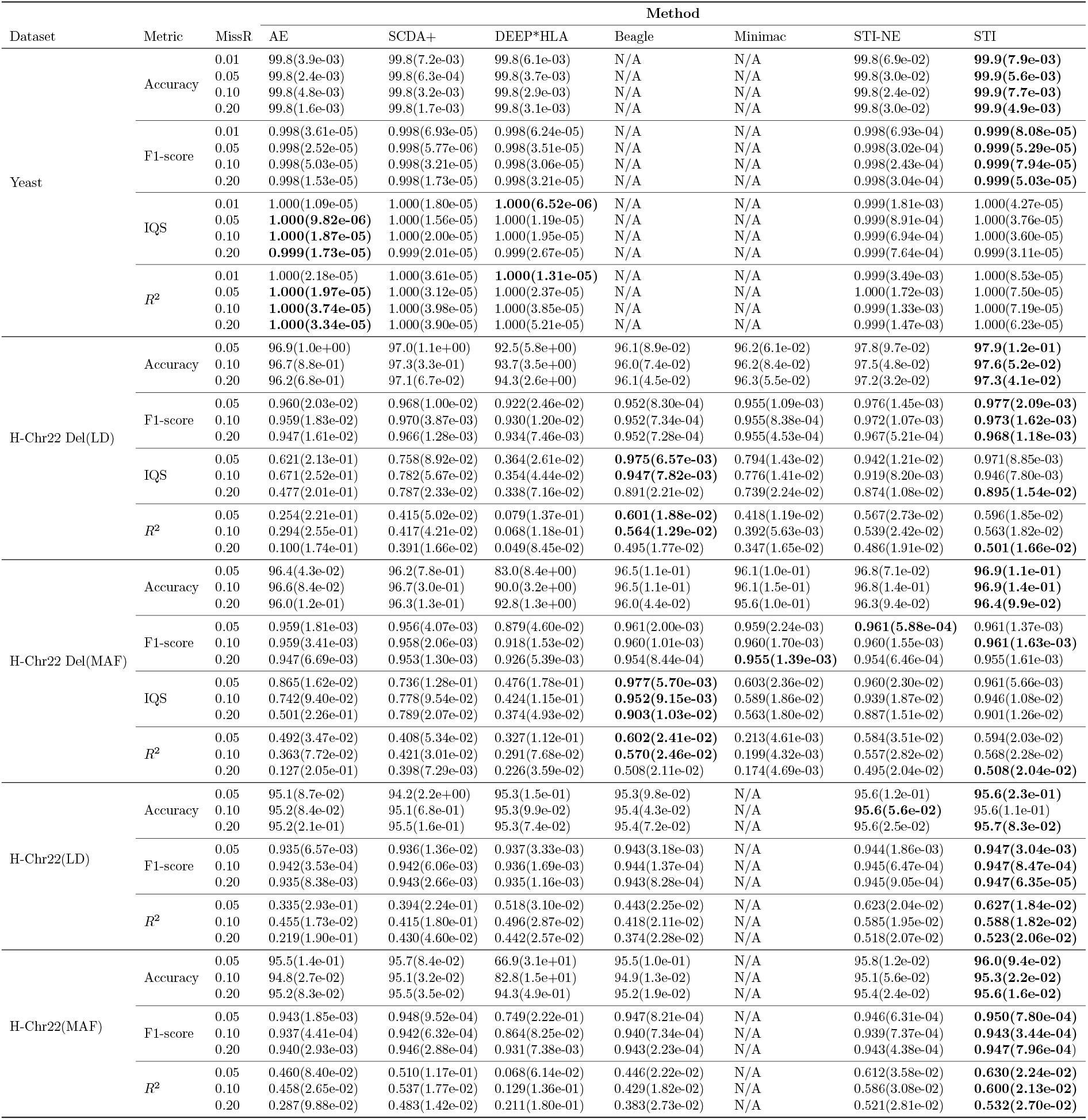
Experimental results for imputing sporadic missingness averaged over 3-fold cross-validation using different MissRs. The numerical values in parentheses show the standard deviation. N/A values indicate that the model could not impute that specific dataset. For all the metrics, the higher the value, the better the models are performing in imputation. In these experiments, accuracy and f1-score calculations distinguish heterozygous alternative alleles by encoding them to distinct categorical values; however, for IQS and *R*^2^ the encoding values remain identical. Bold values indicate the top result in each row. Beagle5.4 generally performs the best in terms of *R*^2^ and IQS for bi-allelic variants, but STI outshines other methods in imputing all SVs (and multi-allelic variants). *Note:* H-in the first column indicates that the dataset is a human genome dataset.

We used maximum LD bins and/or MAF bins (Figure 4 b, and c) to distribute missing positions in the datasets extracted from the human 1000 Genomes Project. If bins had too few positions (e.g., at a 0.01 MissR on chromosome 22 datasets), we excluded this MissR for the experiments related to these datasets. We used a consistent approach to introduce missing values in chromosomes 6, 10, 16, and 22 based on LD distributions and a single test MissR of 0.2. In this experiment, we focused on comparing Minimac, Beagle, and STI, because they were identified as the top performers from classical and DL methods, respectively, in prior experiments. We employed 3-fold cross-validation for both methods, training and imputing each chromosome separately. *R*^2^ was calculated for each variant, and the results were averaged over fold, chromosome, and SV type. Figure 3 presents the experimental results for the extensive structural variation datasets where the top plot shows the improvement that STI provides compared to the best of other methods for each SV type, and is calculated as follows:

**Fig. 3.**
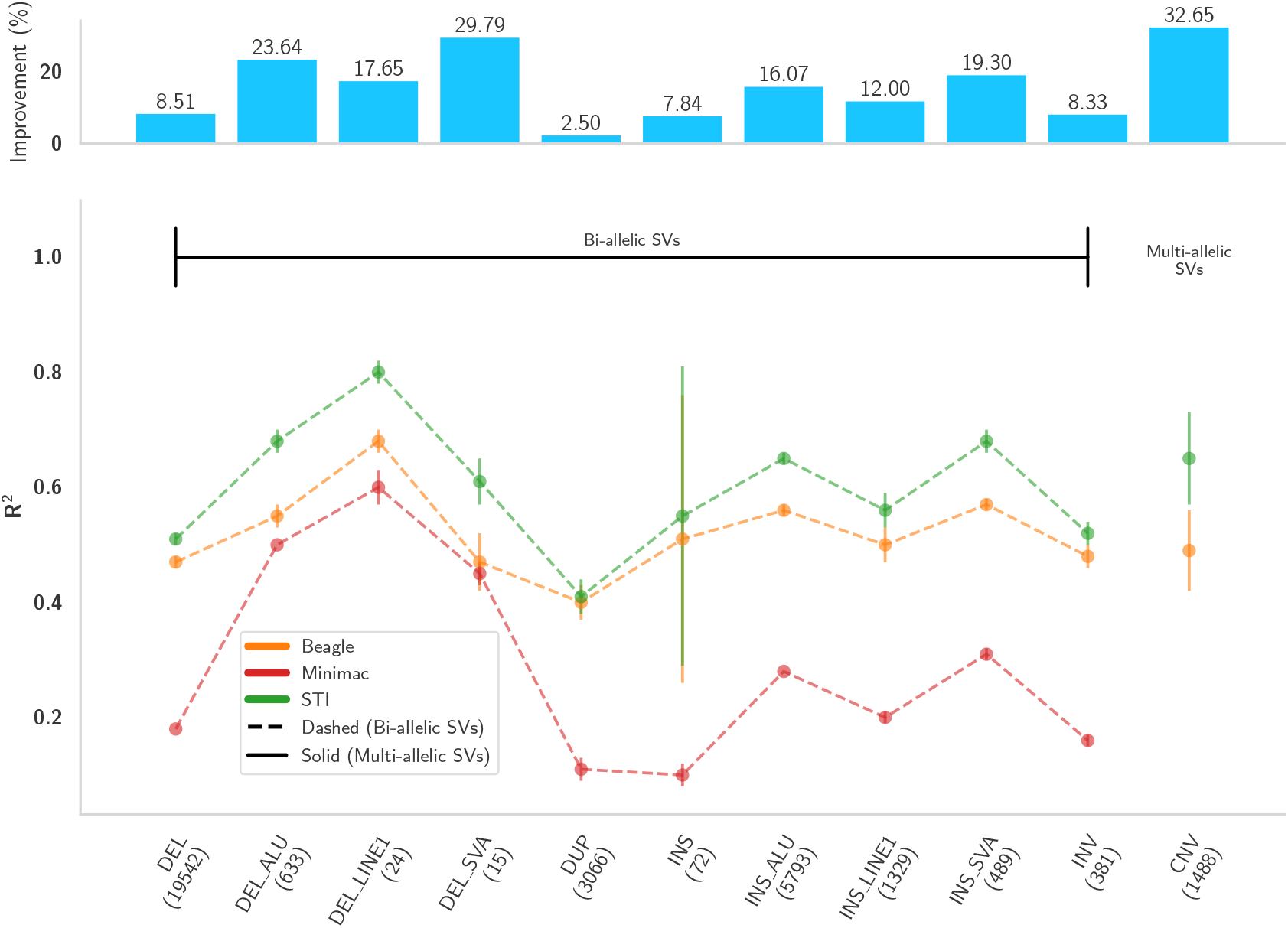
Comparison of Beagle and STI across SV types. Average *R*^2^ of ground-truth genotypes in the test sets and respective predictions over 3-fold cross-validations on chromosomes 6, 10, 16, and 20. The experiments are performed on each chromosome separately, and the results are averaged over chromosomes and folds. Vertical lines indicate standard deviations. The improvement plot shows *R*^2^ score difference between STI and the best of other methods, normalized by the best *R*^2^ scores for each SV type.

**Fig. 4.**
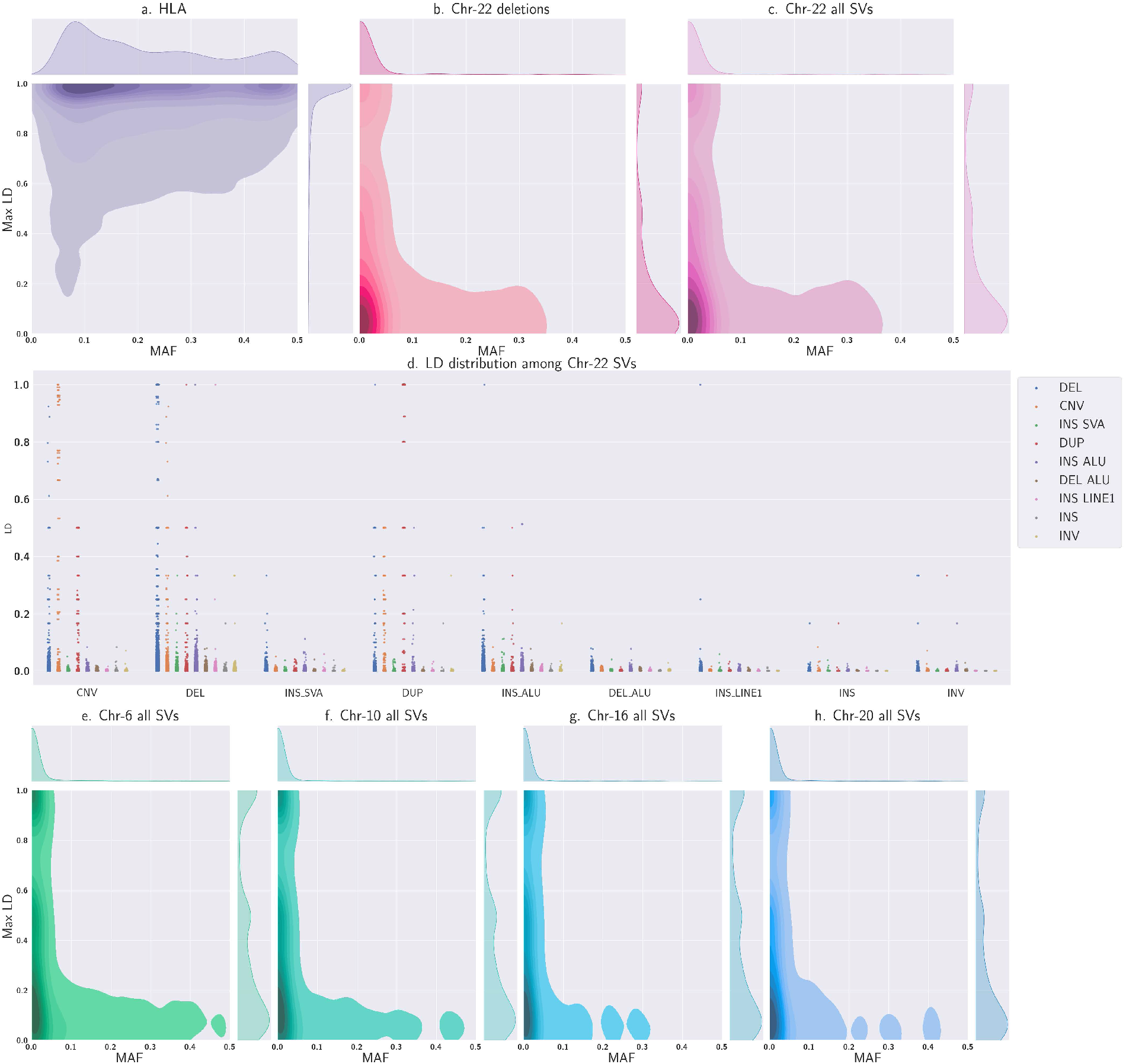
MAF and LD distributions of benchmark datasets from 1000 Genomes Project. MAF and maximum LD distributions are presented using kernel density estimation plots for SNVs/SVs in *a. HLA region on chromosome 6, b. deletions in chromosome 22, c. SVs in chromosome 22, e. SVs in chromosome 6, f. SVs in chromosome 10, g. SVs in chromosome 16*, and *h. SVs in chromosome 20*. Overall, SVs exhibit a low LD value, posing a significant challenge to imputation methods. Plot *d. LD among different SV types in chromosome 22* shows that structural events are commonly correlated with deletions. Furthermore, deletion, copy number variation, and duplication events appear in different ranges of LD, while the rest of the events are limited to *LD* ≤ 0.1. Lastly, the majority of correlated SVs to deletions are of the same event, making deletions a good separate dataset for our experiment.

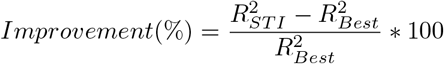

#### Yeast dataset

Missing positions in samples were selected randomly, as the LD analysis showed that the maximum LD for all the SNVs was high in the [0.8, 1.0] range. As mentioned, Minimac4.1.4 and Beagle5.4 cannot be used to impute variants of the yeast dataset due to the lack of a reference panel. However, STI could be applied and outperformed other methods, achieving a minimum average imputation accuracy of 99.86%. Overall, all the applicable models performed well on the yeast dataset, which we attribute to the presence of high LD among SNVs in this dataset.

#### Deletions in chromosome 22

For this dataset, we introduced missing positions proportional to the maximum-LD/MAF distribution Figure 4.b. Overall, STI emerged as the best or the second-best model for imputation across all the metrics. STI was more accurate than others for LD/MAF missingness distribution schema. Furthermore, SCDA+ demonstrates a substantial performance advantage over AE in terms of IQS and *R*^2^ in the majority of the cases. **Table 8** in the Supplement shows the accuracy trends for different maximum LD values for this dataset when missing values are distributed proportional to variant density in maximum LD bins. Minimac4.1.4 and Beagle5.4 were less accurate for SNVs with lower maximum LD compared to AE, SCDA+, STI-NE, and STI. Since HMMs and graphical models rely on conditional probabilities, we suggest that they would perform relatively weak due to a low correlation between the events (states).

#### All SVs in chromosome 22

Similar to the previous dataset, missing positions were distributed among SVs based on maximum-LD/MAF (Figure 4.c). Despite having a reference panel, Minimac4.1.4 cannot be used for some missing variants for this dataset because it can only handle bi-allelic events. Furthermore, IQS is not well-defined for multi-allelic events.

Table 1 shows that STI outperforms all other methods on average accuracy and F1-score. STI performance in terms of *R*^2^ is much better than the competing methods at high MissRs. *R*^2^ considers the correlation among genotypes encoded as categorical values. As such, depending on the difference in encoded values for the predicted and the ground truth genotypes, the penalty can be severe. For example, if 0 | 0, 0 | 1, and 1 | 1 are encoded as 0, 1, and 2 in genotypes and the ground truth for a given genotype is 0 | 0, the model is punished moderately(severely) for predicting 0 | 1(1 | 1). Additionally, SCDA+ outperforms AE in most comparisons, indicating the effectiveness of our proposed training procedure.

#### Extensive structural variation datasets

In this experiment, we focus on *R*^2^ between the predicted and ground truth genotypes as *R*^2^ was the most discriminating metric for comparing the performance in imputing SVs. For estimating *R*^2^, predictions are converted into categorical values, e.g., 0 | 0, 0 | 1, and 1 | 1 are encoded as 0, 1, and 2. Any discrepancy between the model’s prediction and the ground truth leads to a substantial penalty on the correlation, enabling us to see differences more clearly. We found STI consistently outperforms Beagle5.4 and Minimac4 across various SV types, often by a noticeable margin. The underlying cause of this observation is the lack of high LD in this dataset (Figure 4) and fundamental differences between HMMs and the Transformer model. In HMMs, information propagation between two distant variants occurs sequentially through intermediate sites. However, this mechanism falters when the LD block is sparse, leading to reduced performance. In contrast, STI employs a direct variant-to-variant attention mechanism within each chunk without needing to model an intermediate site, which effectively mitigates the limitations posed by a weak LD. Furthermore, the multi-head attention mechanism equips STI to discern higher-order and complex patterns among variants, which appear to be crucial for better imputations in the absence of strong LD patterns. These capabilities highlight STI’s superiority in managing SV imputation challenges where traditional HMM-based approaches may be suboptimal. This is particularly the case for duplications (DUP) and insertions (INS) where STI is able to attain a very high *R*^2^ value. This observation matches our expectations since these two types of SVs are relatively challenging in genotype calling as well [33].

## 3 Discussion

More accurate genotype imputations will improve the performance of downstream functional and biomedical genomic studies. Scientists frequently need to employ multiple tools, adapted based on the degree of missingness and types of variants missing, within individual pipelines to carry out imputations. To address this problem, we have presented STI, a masked DL framework, which appears to be one of the first uses of transformer architecture. While STI is currently limited in few ways, we believe that it is a step towards developing a unified approach for successfully imputing missing values for a range of datasets, from small to large amount of missingness, as well as SNVs and SVs. We explored STI’s performance for a range of masking rates (training) and missing rates (application), which revealed that a single STI model, trained with a masking rate of 0.5, could be applied for imputing SNVs and SVs. STI’s performance in imputing SNVs and SVs was comparable to many other methods and approaches for SNVs and SVs found in low and high LD regions. That is, STI is capable of effectively capturing short and long-range correlations among SNVs/SVs.

STI also performed well in imputing values that were missing systematically (Figure 1.b and Supplementary Information; Ablation and Experimental Results sections). Furthermore, STI offers two additional advantages. First, it can be applied directly to resequencing datasets from any species because, unlike HMMs based on Li and Stephens model [34], a transformer model does not need hard-coded or parametric assumptions about underlying characteristics of the genomic data, such as mutation rate and density of LD blocks, and captures inherent patterns automatically. Secondly, an STI model needs to be trained once to impute sporadic and systematic missingness rapidly and accurately. Therefore, we expect the STI framework to spur the next generation of approaches to advance generalized and efficient imputing, which could served by online imputation servers because of the low computational burden of imputation after training.

Also, STI can be extended by integrating secure and privacy-preserving mechanisms through homomorphic encryption and Tensorflow-compatible libraries such as tf-encrypted [35]) to provide secure and privacy-preserving genotype imputation [36, 37], which is imperative yet under-explored in current tools.

Currently, training an STI model is resource- and time-demanding. One of our development plans for STI is to address these issues. Moreover, STI is a transformer model, and transformers are known to require a large number of samples to achieve optimal performance. Therefore, we expect STI’s performance to greatly improve even for SNVs and SVs in low LD regions as the number of samples is increased due to the release of resources such as those from TopMed and Gnomad4.0.

## 4 Methods

In this section, we first introduce the datasets we used in this study and discuss their characteristics. This is followed by the architectural design of STI and the procedure for model training.

### 4.1 Data

We used five datasets from two sequencing projects [10, 22] to fine-tune and benchmark STI against the baselines. The 1000 Genomes project datasets are pre-phased using Shapeit2. Thus, the phasing information of the test sets is propagated to the training sets, but the process is identical for all the methods so it will not bias the results in favor of any imputation method. Scikit-allel package [38] is employed to compute LD and MAF for the datasets. In the sporadic missingness experiment, we use 3-fold cross-validation to assess the performance of the methods. In a 3-fold cross-validation, each dataset is separated into three distinct partitions where there is no sample overlap. Each time, one of the partitions is used as the test set while the remaining two partitions are used for the training process. For the DL models, a validation set is selected from the training samples for early stopping. To ensure that the same training/validation/test set is used across different methods, we used fixed random seeds for splitting the data into folds and introducing missingness into the test sets.

We used three schemes to introduce sporadic missingness: random selection, MAF-distributed selection, and LD-distributed selection. For the latter two, we computed MAF and LD on the whole data, as depicted in Figure 4, and selected the missing SNVs/SVs for the test set evaluation proportional to those. That is, if 10% of the variants have an MAF in the range of [0.2, 0.3), we selected 10% of the missing values from these specific variants. This approach ensures that the missing data are imputed based on the distribution of MAF or LD in the data, providing a representative imputation strategy. Regardless of the scheme, we used fixed random seeds per sample to decide the missing genotypes. Therefore, the missing genotypes across the test samples are not identical. However, the coordinates of missing values are identical across the competing methods. The characteristics of the datasets we use in our experiments are as follows:

#### 4.1.1 HLA dataset

This dataset contains human leukocyte antigen genotypes, covering a 3 Mbp region at chromosome 6p21.31 and sitting at a major histocompatibility complex (MHC) region. HLA region regulates the immune system in humans [39]. It is highly polymorphic and heterogeneous among individuals; i.e., it harbors various alleles, enabling the adaptive immune system to be fine-tuned [40]. In this study, we used the genotypes of this region, obtained from phase 3 of the 1000 Genomes Project [10], which contained 7161 unique genetic variants for 2504 individuals from five super-populations across the world: American (AMR), East Asian (EAS), European (EUR), South Asian (SAS), and African (AFR). The majority of SNVs in this dataset exhibit maximum LD values in the range of [0.9, 1.0]. We used this dataset for our masking study and fine-tuning the hyper-parameters of DEEP*HLA, SCDA, AE, and STI.

#### 4.1.2 Yeast dataset

The second dataset is the comprehensively assayed yeast dataset [22], representing a simple genetic background and high correlation among genotypes. This dataset contains 4390 genotyped profiles for 28,220 genetic variants. The samples were obtained by sequencing crosses between two strains of yeast, namely an isolate from a vineyard (RM) and a popular laboratory strain (BY). In the original dataset, the data is encoded as -1/1 for BY/RM, which are mapped to 0/1 in our code, respectively, before one-hot encoding.

#### 4.1.3 Chromosome 22 datasets

We used structural variation data from the 1000 Genomes Project in two settings. In the first, we only selected deletions (DEL), excluding ALU/SVA/LINE1 deletions, among all SVs. This resulted in 573 positions harboring bi-allelic events in the dataset. In the second, a total of 848 SVs including, but not limited to deletions, insertions, duplications, inversions (INV), and copy number variations (CNV) in chromosome 22 are selected. As shown in Figure 4 b & c, the majority of SVs in chromosome 22 exhibit a low LD, rendering these datasets challenging for imputation compared to SNVs. According to Figure 4.d, deletions cover a wide range of LD among them and other SVs, making them a good target for a separate bi-allelic dataset.

#### 4.1.4 Extensive structural variation datasets

In the concluding experiment, we undertook a thorough investigation of SV imputation using the human 1000 Genomes Project, selecting 4187, 3126, 2062, and 1569 SVs located in chromosomes 6, 10, 16, and 20, respectively. This selection strategy was informed by the aim to encompass chromosomes of different lengths, providing a representative cross-section of the genome. This diverse chromosome selection allows for a broader understanding of the genomic distribution and characteristics of SVs, facilitating a more nuanced analysis of their presence and impact across different regions of the human genome. These SVs include deletions, duplications, insertions, inversions, and copy number variations. Among these SVs, 469 of them are multi-allelic (CNVs). For each model, we train on and impute each chromosome separately, and take the average of the results over folds, chromosomes, and SV type. The distributions of SVs in these chromosomes in terms of MAF and LD are presented in Figures 4 e, f, g, and h, indicating low LD and diverse MAF in general for the mentioned SV datasets.

### 4.2 Training procedure

In previous studies, the training data is masked using different rates to match the test set, e.g., [3, 14]. In our experiments, we observed improved performance of the model with 50% dynamic and random masking of the variants in the training data. So we trained the DL model once and reused it multiple times. Notably, this masking is similar to the masking performed in modern large language models. The benefit of such a technique in genomic data imputation is the notable reduction in the inference (imputation) times when compared to the fastest traditional methods. Consequently, a DL model trained in this manner becomes particularly advantageous for deployment on imputation servers, where re-training needs to be avoided for quick and efficient processing.

Another improvement we achieved was by representing phased diploids into haploids, followed by one-hot encoding. That is, instead of feeding (one-hot encoded) phased diploids to the models, we fed them haploids. This idea is proposed in [26], but there is no discussion about the merits of this procedure. We surmised that predicting haploids would be easier because mutations in paternal and maternal haploids are independent of each other. In the output, diploid genotypes were reconstructed by combining corresponding haploids together.

### 4.3 Split-Transformer Impute architecture

Split-Transformer Impute is an extended transformer model [28] specifically tailored for genotype imputation. STI models do not require any additional information provided by a reference panel, except for the genotypes and their relative positions. This makes STI adaptable to any genotype data and allows it to be applied to a wider range of datasets with less effort and fewer preparations. Moreover, although here we focus on sporadic missingness, once STI is trained on a dataset, it can predict both sporadic missingness and systematic missingness in genotype data as long as the target variants are a subset of the training variants. An overview of STI is presented in Figure 5. We implemented STI and the rest of the DL models using Tensorflow framework [41] in Python. In order to train the models, we used tensor processing units (TPU) provided by the Google Colaboratory platform, but a GPU implementation of STI is available as well. A learning rate scheduler and early stopping are employed in order to reduce the loss and training duration.

**Fig. 5.**
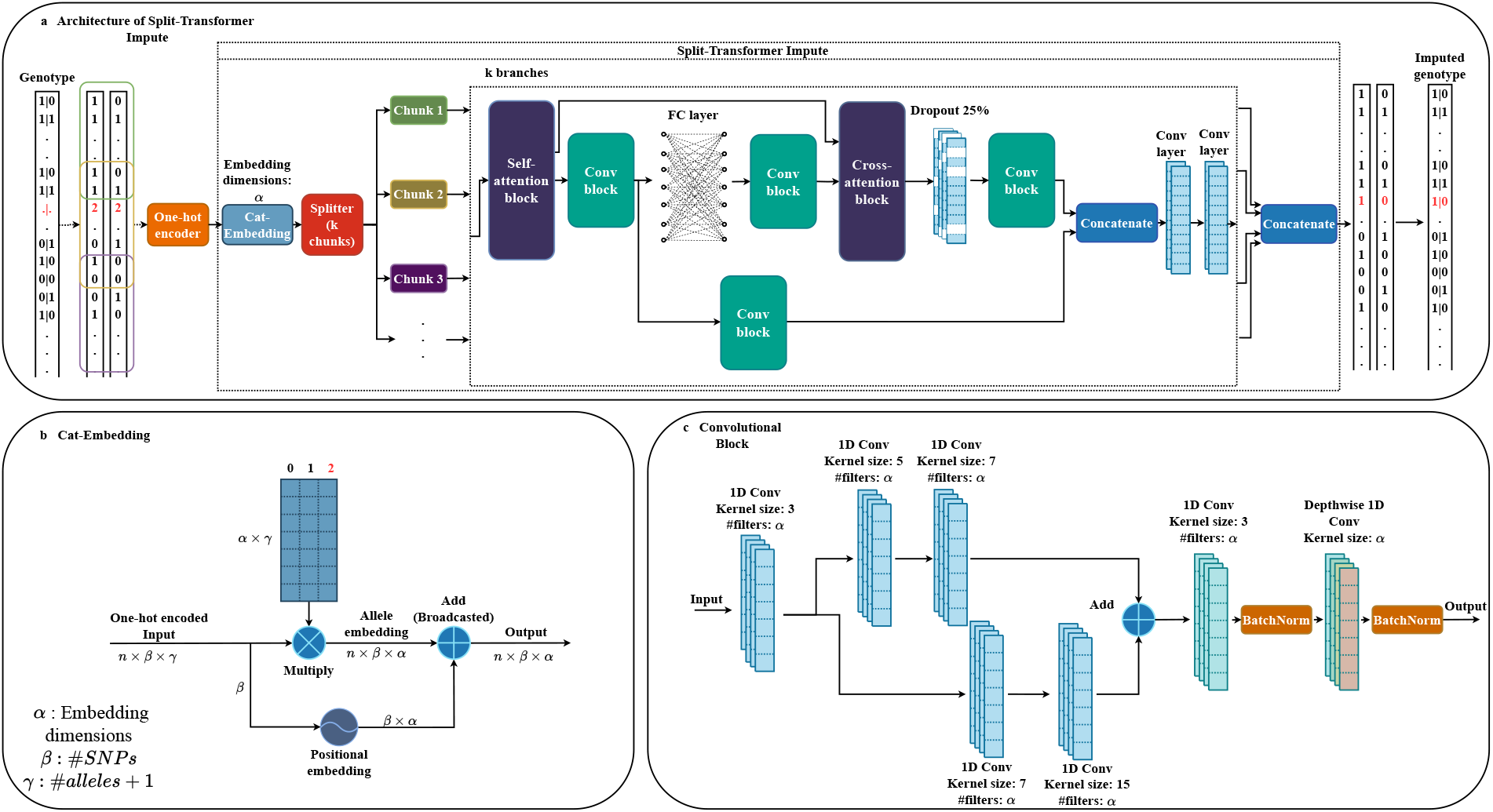
Split-Transformer Impute architecture. *a. Overall pipeline of the proposed framework:* the data is separated into paternal and maternal haplotypes in the case of diplotypes, and it remains the same for haplotypes. While the figure shows phased genotypes, STI can handle unphased data as well (though the performance degrades). Next, the data is one-hot encoded and fed into our Cat-Embedding layer, followed by splitting the data vertically into k chunks. The chunks have overlap in order to capture information for the SNVs residing around the chunks’ edges. Each branch passes through a unique set of attention, convolution, and fully connected layers. In the self-attention block, the flanking variants that come from the neighboring chunks are removed after applying multi-head attention. Finally, the results of all branches are assembled to generate the final sequence. *b. Workflow of proposed Cat-Embedding:* we consider a unique vector space for each unique categorical value in each SNV/feature. To save computational resources, instead of pre-allocating these vectors, we use the addition of positional embedding and categorical value embeddings in order to generate unique embedding vectors for each categorical value in each SNV/feature. We consider a missing (or masked) value as another categorical value (allele) in our model. Here, 2 (highlighted in red) represents the missing value. *c. Convolution blocks:* two parallel convolutional branches with varying kernel sizes are used in our convolution blocks. These multi-scale convolutional blocks allow STI to capture information at multiple spatial scales in the input data, similar to the pattern-matching idea used in classical computer vision methods using convolution. Given the variable sizes of LD blocks, multi-scaled convolution is expected to excel at capturing LD patterns compared to single-scaled convolutions.

#### 4.3.1 Cat-Embedding

One important part of STI is categorical embedding (Figure 5.b), termed as Cat-Embedding, which enables it to learn embedding representation per allele in each position. For the imputation task, we consider missing values as another allele that is equivalent to special tokens in natural language processing. The corresponding vector for each allele is added to the respective positional variant embedding vector to generate the final embedding. The idea is similar to a natural language processing embedding layer that accepts word indices, except that Cat-Embedding accepts one-hot encoded data.

#### 4.3.2 Splitting

While the multi-headed attention in a transformer offers significant advantages, a major drawback is quadratic memory cost for computations that becomes important in genomic analysis, since the number of variants in a sample is normally in the thousands. In genotypes, the majority of interactions are local [42]. Therefore, it is of great importance to limit the scope of attention to save computational resources. To do so, we split the variants into chunks (vertical partitioning). The chunk size and overlap size are employed in a comparable manner in Minimac4.1.4 and analogous software applications. In order to prevent loss of imputation accuracy at chunk borders, we include flanking variants from neighboring chunks and discard them after applying self-attention to get the original variants in the chunk. Though the average LD block size in the dataset can be used to decide the size of overlap, we do not use LD blocks directly to decide the chunk size in the current version.

Each chunk passes through a dedicated branch inside the model, leading to increased imputation quality. Ideally, having a vast number of samples allows training a single model with attention across the whole genome. However, when the number of samples is not enough, the model is left with untrained parameters, resulting in poor performance. Hence, chunking regulates the number of parameters. In a vanilla transformer, the cost of computing global attention is quadratic with respect to the number of SNVs (*m*^2^); however, the amount is lowered to (*m/w*) × (*w* + *o*)^2^ = *mw* in STI, considering that the overlaps of chunks are negligible compared to the chunk size. For instance, for *m* = 10^4^ and a chunk size of 10^3^, STI uses 10 times less memory for attention computations compared to a vanilla transformer.

#### 4.3.3 Attention

The attention blocks are implemented similarly to those of other transformers, such as self-attention blocks in Vision Transformer (ViT) [43]. There is a difference between the first and second attention blocks in the branches. The first block is a self-attention block, meaning that the query, key, and value of the attention layer are the same. The output of multi-head attention in Tensorflow has the same dimensions as the query. By excluding the neighboring variants of a chunk from the query and only including them in the key and value, we involve them in the attention mechanism and, at the same time, shrink the output of a chunk to the target size (chunk size without counting flanking/overlap variants) after applying multi-headed attention. In the second block, the query is the output of the previous layer, while the key and value are the outputs of the first self-attention block. This skip connection considerably affects the overall performance of the model.

#### 4.3.4 Convolutional block

Convolutional blocks, as illustrated in Figure 5.c, are also crucial components of STI. Through empirical studies, we found that using two parallel convolutional branches with varying kernel sizes, similar to the Inception module [44], is the best trade-off between accuracy gain and increase in a number of model parameters, compared to using a single branch or more than two branches. Furthermore, a Depth-wise convolutional layer at the end of the block helps STI extract local information without mixing channel information and substantially improves imputation accuracy.

#### 4.3.5 Output formation

Finally, the outputs of all branches are concatenated to form the output, that is, either maternal or paternal haplotype in the case of 1000 Genomes Project datasets or the genotypes in the case of yeast. For the former, by assembling maternal and paternal haplotypes, we obtain imputed genotypes, and the latter needs no further post-processing. Since genetic variations in parents are independent, directly encoding and imputing the genotypes in diploid life forms results in lower imputation accuracy compared to imputing their haplotypes. Hence we undergo extra steps to separate diplotypes into haplotypes in pre-processing, and combining respective predicted haplotypes into diplotypes in post-processing for the human 1000 Genomes Project dataset.

#### 4.3.6 Loss function

For the loss function, we used a combination of Kullback–Leibler divergence (*D*_*KL*_) and categorical cross-entropy (CCE), similar to the loss function of variational autoencoder [45], as follows:

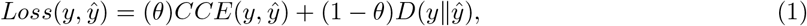

where *θ* is the weight parameter. The first term, representing categorical cross-entropy, and the second term, representing Kullback–Leibler divergence loss, are calculated as follows:

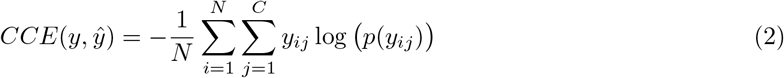

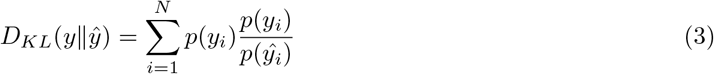

We set *θ* to 0.5, meaning that STI minimizes Equations 2, 3 equally. CCE captures reconstruction error between the input and the output, while *D*_*KL*_ measures asymmetric distance, with *y* as the base, between their probability distributions. In our experiments, omitting any of these losses resulted in reduced model performance. Theoretically, KL-divergence and cross-entropy are related and using both might not seem to contribute to the performance of the model. However, adding KL-divergence to the loss term helps the model to retain the probability/dosage distribution of alleles per variant. In other words, while crossentropy focuses on predicting the correct genotype, KL-divergence acts as a regularization factor and penalizes the model whenever the shape of the predicted probability distribution (allele probability/-dosage) shows divergence from the ground truth. Moreover, the mathematical relation of *D*_*KL*_ and *CCE* can be summarized as follows:

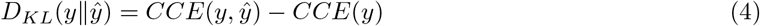

where *CCE*(*y*) is the entropy of the ground truth. According to Equation 4 minimizing *D*_*KL*_(*y* ∥ *ŷ*) is equivalent to minimizing *CCE*(*y, ŷ*) under the condition that the entropy of the ground truth remains constant. However, in deep learning models, data is typically processed in mini-batches. This means that the entropy of each mini-batch may not accurately represent the entropy of the entire ground truth. As a result, Equation 4 does not hold for the DL models in general.

### 4.4 Baseline models

In order to benchmark our model, we compare STI to state-of-the-art imputation models capable of imputing sporadic missingness: SCDA [14], AE [3], DEEP*HLA [26], Beagle5.4 [19], and Minimac4.1.4 [15]. In [14], experimental results indicate that SCDA outperforms classical ML models for genotype imputation. Hence, we do not include classical ML models in our benchmarking analyses. It is worth noting that DEEP*HLA is not originally designed for genotype imputation in general, but we modified and fine-tuned it to work for this problem. Additionally, in order to assess the contribution of Cat-Embedding, we replaced it with a convolution layer in STI, named the resulting model STI-NE, fine-tuned it, and applied it to the benchmark datasets. Lastly, we trained SCDA, in addition to DEEP*HLA and STI, using our proposed pre-processing and training procedure, and compared it to AE. Since AE and original SCDA are the same and only differ in the training process (which results in AE outperforming SCDA), we believe that this comparison can demonstrate the effectiveness of our proposed pre-processing and training procedure.

For SCDA and AE, hyper-parameter tuning information on the yeast dataset is present in the original papers. For SCDA, AE, DEEP*HLA, and STI, we conducted a grid search for optimal hyper-parameters on the HLA dataset using validation sets in a 3-fold cross-validation. We assessed the impact of these hyper-parameters on the performance of the models within the HLA dataset and applied these findings to select suitable hyper-parameters for the yeast dataset in the case of DEEP*HLA and STI, and for the SV dataset across all four mentioned methods. The upper limit for the hyper-parameters was the resource limit of Google Colaboratory using Nvidia Titan IV GPU with 16 GB of RAM size for AE, and roughly the same limitation for TPU RAM size. Minimac4.1.4 and Beagle5.4 do not require fine tuning for the experiments we run.

### 4.5 Experimental settings

The input to all DL models is one-hot encoded. While STI can handle diploids, we found that the best performance was achieved when the inputs of the DL models were haplotypes, an analysis inspired by [26]. Therefore, for the HLA dataset and chromosome 22 datasets, we separated each diplotype into maternal and paternal haplotypes, fed them into the model, and reconstituted the resulting predictions for DEEP*HLA [26], SCDA [14], and STI. We continue using diplotypes as inputs for AE [3] since it is an improved version of SCDA in which the training process was modified, and we wanted to keep it intact. By doing so, we also compare the improvement in AE to our implementation of SCDA, called SCDA+, in which we use proposed pre-processing in conjunction with the changes to the training process as a contribution. The yeast dataset contains haplotypes, so there is no need for the aforementioned extra steps.

In this study, to evaluate the imputation power of the models, multiple evaluation metrics are used including imputation accuracy, imputation quality score (IQS) [46], weighted *F* 1-score, and correlation between imputed and real genotypes in terms of *R*^2^ [47]. Accuracy and weighted *F* 1-score are calculated only for positions with missing genotypes and for these metrics, heterozygous genotypes are encoded differently; i.e., *0* | *1* and *1* | *0* are encoded to two different categorical values. IQS adjusts the chance concordance between predicted and the ground truth SNVs and is defined for bi-allelic events. Therefore, IQS cannot be calculated for any SV in chromosome 22. *R*^2^ is the squared Pearson correlation coefficient between the imputed genotypes and the true genotypes at a specific locus. The definition of these metrics is provided in the Metrics section of the Supplement.

## Data availability

All data used in this study are publicly available. The yeast dataset can be found as the *Supplementary Data 5* at https://www.nature.com/articles/ncomms9712, the rest of datasets are extracted from the 1000 Genomes Project phase 3 dataset available at http://ftp.1000genomes.ebi.ac.uk/vol1/ftp/release/20130502/. Instructions on how to prepare the data for Missing variants experiment can be found in https://github.com/kanamekojima/rnnimp.

## Code availability

The source code of STI is publicly available on GitHub (https://github.com/shilab/STI).

## Acknowledgements

This work is partially supported by the US National Science Foundation (DBI 1750632) and the National Institutes of Health (GM-0126567-03). This research includes calculations carried out on HPC resources supported in part by the National Science Foundation through major research instrumentation grant number 1625061 and by the US Army Research Laboratory under contract number W911NF-16-2-0189. We appreciate the suggestion provided by Dr. Francisco McGee in designing the model, leading to improvements in performance. Additionally, we would like to thank Dr. Kaname Kojima for helping us obtain the data for the Missing variant experiment and providing experimental results for RNN-IMP (the experiment in the supplement) and Emily Thyrum for proofreading the manuscript.

## 5 Contributions

M.E.M. developed the method with the help of J.C., B.J., V.C, and X.S. and M.E.M. implemented the code. C.L. and O.J. prepared the datasets. M.E.M. and R.D. performed the experiments. M.E.M, S.K., C.L., O.J., and X.S. conducted the data analysis. M.E.M, C.L., S.K., T.R.R., V.C, and X.S. wrote the manuscript. All the authors read and approved the submitted manuscript.

## Ethics declarations

Not applicable.

## Declarations

Not applicable.

