## Supplementary information for "Split-Transformer Impute (STI): A Transformer Framework for Genotype Imputation"

#### 1 Introduction

Previously, in the main manuscript, we discussed the importance and applications of imputations in recent years. However, imputation is but one approach to handle missing value. In this section, we discuss the alternative approaches in order to provide a more inclusive background for the problem.

##### 1.1 General approaches to handle missing values

Faced with missing values in the data, there are two general approaches we can follow, ignoring missing values and data imputation.

*Ignoring Missing Values*: If the number of samples containing missing values are close to nothing and we have abundant samples, it is a naive yet common practice to just remove those samples [1]. Alternatively, we can remove the features that are not informative and contain missing values. A better approach is to keep all samples but use non-missing features per sample in analysis so that we do not discard potentially useful information. The drawback of this approach is that it uses varying sample size per feature [2].

*Missing Value Imputation:* We can use computational methods to infer the original value for missing values in the data, otherwise known as data imputation [3]. Imputation methods in general fall in either single imputation (SI) methods or multiple imputation (MI) methods. SI methods replace missing values by a single value, e.g., the mean of the values for that feature [4]. In a special case that the feature/variable is categorical, we can put a special value to represent missing values and let the learning algorithm handle the missing parts, though the performance is expected to deteriorate since the model is missing the information [5]. The drawback of replacing by a constant is that it may change the distribution of the data and introduce bias [6]. An alternative and more sophisticated method is to regress the missing values. In this method, missing value is replaced by predicted data using prediction algorithms such as K-Nearest Neighbours (KNN) [7, 8], Hidden Markov Models (HMM) [9, 10], and Deep Learning (DL) models [11, 12]. In MI, for each missing value, multiple possible values are generated with varying probabilities to account for uncertainty of predicted values [13, 14]. As a result, MI methods generate m different datasets at the end of their process [5].

#### 2 Ablation study

To decide which architecture is the best choice for STI, we performed an ablation study using three scenarios. In the first, we removed one branch in convolutional blocks (Figure 1.c, bottom branch). In the second scenario, we remove convolutional blocks and the dense layer in the middle of the branches entirely and just used transformer blocks in the models, with the exception of the output layer in each branch pertaining to a chunk. Lastly, we removed the transformer blocks to monitor how the model performs without any attention. We used a 3-fold cross validation on the HLA dataset and optimal hyper-parameters we found for STI during hyper-parameter tuning for all these models. Missing values were introduced to the data proportional to Minor Allele Frequency (MAF) distribution, in a similar fashion to the sporadic missingness experiments in the main manuscript. The experimental results of this study are available in Table 1.

**Table 1** Ablation study results for alternative architectures of STI, using a 3-fold cross validation on HLA dataset. STI-NA is STI without any attention blocks. STI-1BC is the case that we remove one of the branches (the branch with larger kernel sizes) in convolutional blocks in STI. 2T, 4T, and 8T are the variations of STI in which convolution is removed and there are respectively 2, 4, and 8 transformer blocks in the model. Numerical values in each cell show the accuracy and the values in parentheses indicate standard deviation. Bold values are the top results in each row.

| | Method | | | | | |
|---|---|---|---|---|---|---|
| Test MissR | STI | STI-NA | STI-1BC | STI-2T | STI-4T | STI-8T |
| 0.01 | **99.2(3.1e-02)** | 99.1(1.2e-01) | 99.0(5.7e-02) | 99.0(3.2e-02) | 99.1(5.0e-02) | 99.0(6.4e-02) |
| 0.05 | **99.2(4.7e-03)** | 99.1(7.3e-02) | 99.1(4.5e-02) | 99.0(6.3e-04) | 99.0(3.3e-02) | 99.0(4.6e-02) |
| 0.10 | **99.1(2.3e-03)** | 99.1(6.9e-02) | 99.0(2.5e-02) | 99.0(1.0e-02) | 99.0(3.6e-02) | 99.0(4.6e-02) |
| 0.20 | **99.1(1.3e-02)** | 99.0(6.0e-02) | 99.0(2.6e-02) | 98.9(1.2e-02) | 98.9(4.0e-02) | 98.9(6.3e-02) |

Observations suggest that attention (STI compared to STI-NA) might not significantly contribute to the performance. However, we suspected that the reason attention is not outshining convolution here is the fact that many variants in this dataset have a high Linkage Disequilibrium (LD) and MissR in the test set is not high enough for the transformer to make a considerable difference, i.e., when the MissR is low, the chance of finding a neighboring variant that is in high LD with a missing variant is high.

Therefore, we used the SNVs in chromosome 22 dataset from human 1000 genomes project, and used PLINK2 [15] to remove missing values, multi-allelic events, and the variants which exhibit a Hardy-Weinberg equilibrium test $p$-value below 0.01, resulting in a total of 31143 variants for the reference panel. Following the instructions for data collection in [16] and a script provided by the authors, we created a microarray dataset using Infinium Omni 2.5 BeadChip manifest for the same chromosome with 7336 variants. We followed [16] for selecting the exact individuals for the microarray data (test set), and the rest for the reference panel. Then we trained STI and STI-NA using MaskR=0.8 on this reference panel of chromosome 22 dataset, and imputed Omni2.5 microarray genotype data. The results of this experiment is available in Table 2. As surmised, STI outperformed STI-NA with a higher performance gap compared to the previous experiment. Notably, this performance discrepancy was more pronounced in variants within weak LD blocks. This finding supports our earlier assumption about the negligible performance difference between the two models SNVs in strong LD blocks.

**Table 2** Ablation study results obtained from training the models on WGS chromosome-22 dataset and imputing systematic missingness in Omni2.5 microarray data. The values show the accuracy of the methods. Both are trained using MaskR=0.8.

| | LD | | | | | |
|---|---|---|---|---|---|---|
| Method | [0, 0.2) | [0.2, 0.4) | [0.4, 0.6) | [0.6, 0.8) | [0.8, 1) | 1 |
| STI | 85.28 | 88.16 | 91.78 | 93.97 | 97.67 | 99.78 |
| STI-NA | 83.41 | 85.71 | 89.59 | 92.09 | 96.64 | 99.50 |

### 3 Hyper-parameter tuning

In this section we provide the details of hyper-parameter tuning (HPT) performed for the benchmark models and our proposed model. Notably, every metric reported in this section is over all predictions, including masked and non-masked genotypes.

For hyper-parameter tuning using the yeast and HLA dataset, we used 3-fold cross validation and averaged the results over the fold per set of hyper-parameters. Missing values are distributed randomly for the yeast dataset and randomly proportional to MAF distribution for the HLA dataset in validation sets.

STI has four hyper-parameters, namely chunk size, chunk overlap size, embedding dimension ($\alpha$), and number of attention heads. We only tuned STI (and STI-NE) using the validation results of HLA dataset and used the similar settings for other datasets. Based on the validation results in Figure 1 and Table 5, we observe that increasing the number of attention heads and chunk size, in general, can positively affect the performance of STI, while increasing $\alpha$ and overlap size does not necessarily improve the performance. Hence, in the case of chromosome 22 datasets where we are not hitting memory limitation, we increase the number of attention heads to 40 for actual training and evaluation. The details of hyper-parameter tuning for STI is provided in Figure 1 and Tables 3 & 4 for the HLA dataset. Moreover, we fine-tuned STI-NE only on $\alpha$, and for the rest of STI-NE hyper-parameters we used the best hyper-parameters of STI. In this case, an increase in $\alpha$ was not consistently improving the performance metrics of the validation set. The details for hyper-parameter tuning of the HLA dataset is provided in Table 5.

In case of SCDA and AE, we use the same hyper-parameters reported in the papers for the yeast dataset, but we performed hyper-parameter tuning on the HLA dataset and used the observed results to select proper hyper-parameters for the benchmark datasets. To do so, we use the reported optimal regularization coefficient of the yeast dataset for SCDA and AE in every experiment. Furthermore, we only used SCDA to tune the model since AE differs from SCDA only in training procedure and the architectures are the same. The results of SCDA hyper-parameter tuning are presented in Table 6. Similar to STI, we use optimal hyper-parameters from HLA dataset for chromosome 22 datasets.

We re-implemented DEEP*HLA in Tensorflow, with a minor change to make it suitable for imputing the genotypes in a variant-level rather than gene-level, and fine-tuned it on yeast (Table 7) and HLA (Figure 2) datasets, using a range of filter and kernel sizes for convolution layers. For chromosome 22 datasets, we use the best hyper-parameters of HLA dataset for each model but whenever we were not hitting resource limitations, we performed minimal changes in hyper-parameters for each model to get

**Table 3** Hyper-parameter tuning details of STI using HLA dataset on validation sets. Best results in each block are in bold typeface, while the next best result in in italic typeface.

| Embed dim | Chunk size | Num heads | Max accuracy |
|---|---|---|---|
| 64 | 500 | 16 | 99.625 |
| | | 24 | 99.599 |
| | | 32 | 99.610 |
| | | *40* | *99.628* |
| | **1500** | **16** | **99.635** |
| | | 24 | 99.632 |
| | | 32 | 99.617 |
| | | 40 | 99.627 |
| | 3000 | 16 | 99.607 |
| | | *24* | *99.619* |
| | | 32 | 99.591 |
| | | 40 | 99.612 |
| 128 | 500 | 16 | 98.302 |
| | | *24* | *99.658* |
| | | 32 | 98.687 |
| | | 40 | 99.478 |
| | **1500** | **16** | **99.670** |
| | | 24 | 99.657 |
| | | 32 | 99.660 |
| | | 40 | 99.666 |
| | 3000 | 16 | 99.666 |
| | | 24 | 99.667 |
| | | 32 | 99.665 |
| | | *40* | *99.668* |
| 256 | 500 | *16* | *99.644* |
| | | 24 | 99.633 |
| | 1500 | *16* | *99.678* |
| | | 24 | 99.675 |
| | **3000** | 16 | 99.682 |
| | | **24** | **99.684** |

better results. For instance, we increased the number of attention heads in STI, when the resource limit allows us, since hyper-parameter tuning showed that an increase to the number of heads has a non-negative effect on the performance.

### 4 Experimental results

We presented the key experimental results in the main material and here we will provide additional details.

Accuracy breakdown for deletions in chromosome 22 is available in Table 8. According to these results, STI is generally outperforming competing methods, specially for SNVs in low LD blocks. Moreover, we can observe that there is a trend of increasing accuracy with an increase in maximum LD, except for the last bin ($[0.8, 1]$) where there is a considerable drop in accuracy. We could not explain this drop with preliminary exploratory data analysis and there is a need for further analysis to uncover the underlying cause for this phenomena.

Moreover, to shed light on the performance of the models over different SVs in the test sets of chromosome 22, we extracted every position used to implant missing values and summarized the analytics of these positions and respected performance of competing models in Table 9. The overall performance of the models is competitive in most of events, while STI is outperforming the other methods in most of the cases. Among all SVs, CNVs are multi-allelic variants and STI is outperforming the rest with a considerable accuracy gap.

Though STI excels at structural variation imputation, it is designed to be able to impute SNVs as well. To assess this, we used whole-genome sequencing SNVs in chromosome 22 as the reference panel and Infinium Omni 2.5 Beadchip manifest to extract microarray genotypes, as previously described in

**Table 4** Hyper-parameter tuning details of STI using HLA dataset on validation sets for the chunk overlap size using $\alpha$ of 256 and 24 attention heads. Best results in each block are in bold typeface, while the next best result in in italic typeface.

| Chunk size | Chunk overlap | Max Accuracy |
|---|---|---|
| 1500 | 0 | 99.677 |
| | 50 | 99.678 |
| | 100 | **99.680** |
| | 200 | *99.679* |
| | 400 | *99.679* |
| 3000 | 0 | 99.674 |
| | 50 | *99.680* |
| | 100 | **99.684** |
| | 200 | *99.680* |
| | 400 | 99.671 |

**Table 5** Hyper-parameter tuning results of STI-NE using HLA dataset on validation sets for $\alpha$ using chunk size of 3000, chunk overlap size of 100, and 24 attention heads. Best result is in bold typeface.

| Embed dim | Max Accuracy |
|---|---|
| 64 | 98.909 |
| 128 | **99.423** |
| 256 | 79.101 |

the ablation study. For this experiment, we trained STI on Temple University's HPC server using A100 GPUs. We asked the authors of RapidAE[17] RNN-IMP[16] to impute the data using their models and imputed the microarray data using Minimac and Beagle. The results of this experiment is presented in Table10. As a side note, since GRUD [18] performs equally as good as RNN-IMP, we did not include GRUD in this experiment. RNN-IMP enjoys the highest accuracy among DL models, however it needs to be re-trained each time the set of training/target SNVs change, making it hardly applicable to real world scenarios. Nevertheless, among these deep learning methods, STI is exclusively capable of handling multi-allelic events, while its performance is competitive to Minimac4.1.2 by a margin of 0.5% in terms of accuracy. Since transformer models are known to require more data samples compared to other DL models, we expect that using larger datasets in the next studies can close this performance gap.

Lastly, we present the inference time of competing methods in Table 11. Beagle and Minimac are executed on Temple University's HPC server using 88 Intel Xeon (Cascade Lake) CPU cores. Beagle employed maximum number of threads (176) while Minimac utilized 10 threads. DL models are all evaluated on Google Colab's TPUs, excluding AE. However, the performance of Nvidia A100 GPUs are quite similar to a TPU resource allocated to Google colab users, hence the runtime of AE and SCDA+ are the same.

### 5 Evaluation metrics

We used accuracy, imputation quality score, weighted f1-score, and $R^2$ as the metrics for our evaluations. Accuracy is generally defined as follows:

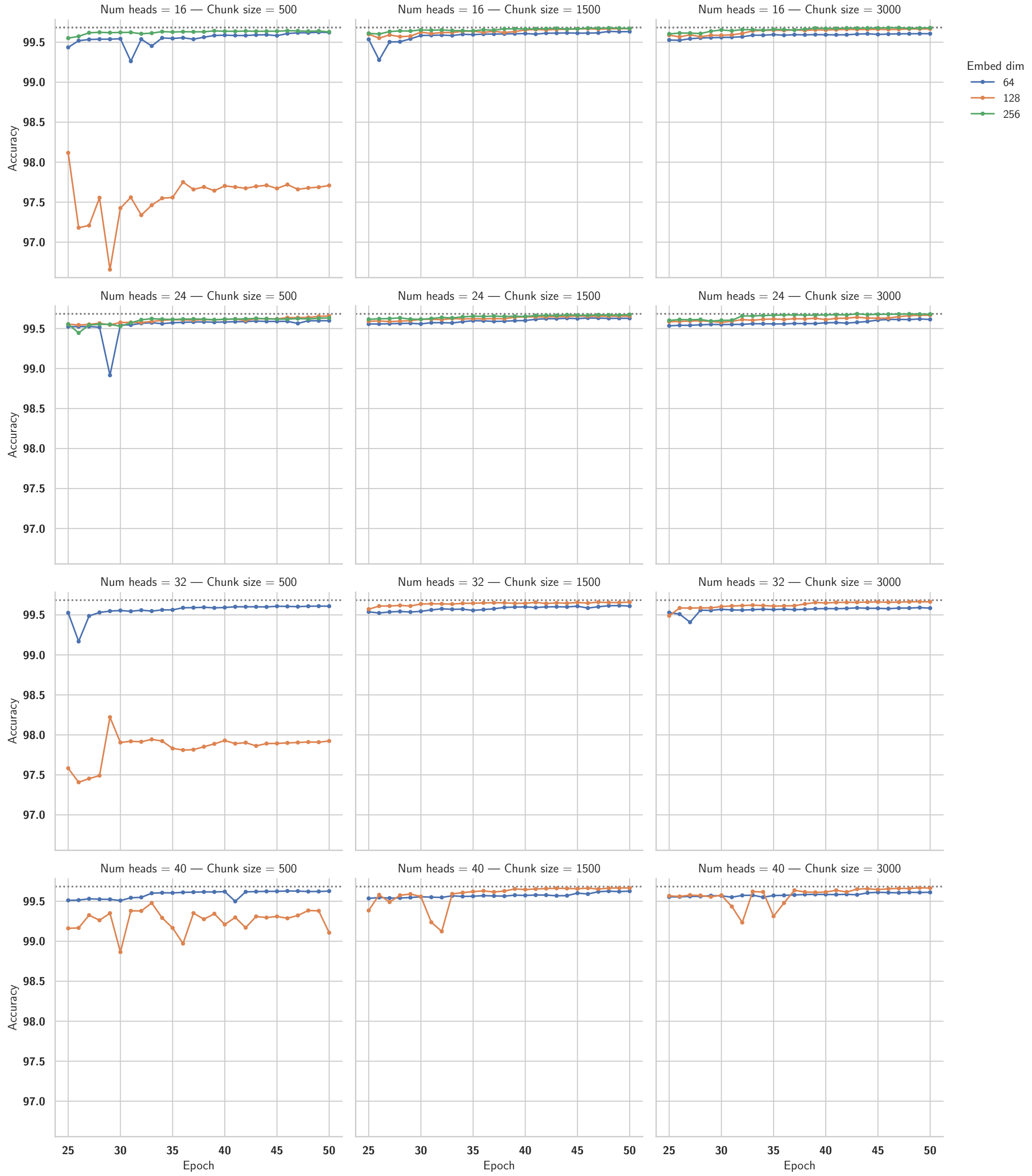


**Fig. 1** The effect of different hyper-parameters of STI on average accuracy of validation sets in 3-fold cross validation on the HLA dataset, where the MissR is 50%. We investigated the impact of embedding dimensions (Embed dim), number of attention heads, and chunk size in STI performance for HLA dataset. Based on these results, embedding dimension of 256 performs the best while increasing the number of heads and chunk size marginally improves the performance. Since accuracy here does not exclusively reflect missing SNVs but also non-missing SNVs as well, any marginal improvement will be significant for actual missing positions.

$$Accuracy = \frac{CP}{N} \tag{1}$$

in which CP and N refer to correct predictions and all possibilities respectively. This term is referred to as concordance rate (CR) in [19, 20]. Simply put, for this problem, accuracy is the percentage of correctly imputed SNVs/SVs among all possible SNVs/SVs. The accuracy can be reported on all the SNVs/SVs in the test set, or only on the missing positions. In our case, we use the latter to report the results.

While accuracy is a good metric to evaluate the performance of imputation models, it does not reflect the performance of the model with regards to hard-to-impute SNVs. IQS differentiates well-imputed and poorly imputed SNVs [21], and is calculated as follows:

**Table 6** Hyper-parameter tuning results of SCDA (and AE) using HLA dataset on validation sets. #Base filters and kernel size refer to filter size for the first convolution layer and size of the kernel for all layers in the models. Best results are presented in bold.

| #Base filters | Kernel size | Max Accuracy |
|---|---|---|
| 64 | 16 | 96.848 |
| | 32 | 98.430 |
| | 64 | **98.611** |
| | 128 | 98.534 |
| 128 | 32 | **98.933** |
| | 64 | 98.822 |
| | 128 | 98.245 |
| | 256 | 96.322 |
| 256 | 64 | **98.605** |
| | 128 | 97.033 |
| | 256 | 96.438 |

**Table 7** Hyper-parameter tuning results of DEEP*HLA using Yeast dataset on validation sets. #Filters 1/2 refer to filter sizes for the first/second convolution layers and kernel size refers to the size of kernel for all layers in the models. In this hyper-parameter tuning, we use only one branch in the DEEP*HLA, whereas in the main experiments we break the input into fix-sized chunks, similar to original DEEP*HLA. Best results are presented in bold.

| #Filters 1 | #Filters 2 | Kernel size | Max Accuracy |
|---|---|---|---|
| 128 | 64 | 64 | 91.649 |
| 256 | 128 | 128 | 95.704 |
| 512 | 256 | 256 | 97.515 |
| 1024 | 512 | 512 | **97.906** |

$$IQS = \frac{P_o - P_c}{1 - P_c} \quad (2)$$

where $P_o$ and $P_c$ are observed proportion of agreement and chance of agreement, and defined as follows:

$$P_o = \frac{\sum_i n_{ii}}{n_{..}} \quad (3)$$

$$P_c = \frac{\sum_i n_{i.} n_{.i}}{n_{..}} \quad (4)$$

in which $n_{i.}$ and $n_{.i}$ are marginal frequencies, meaning that they occur if genotypes are called randomly at the same rate. In other terms, if $a_1$, $a_2$, and $a_3$ are probabilities of AA, AB, and BB for the ground truth respectively and $b_1$, $b_2$, and $b_3$ are respective probabilities of the same based on the imputation results, $n_{ij} = a_i b_j$. IQS accounts for allele frequency by subtracting the chance of agreement from the observed agreement [21].

F1-score is a balanced score reflecting the performance in terms of both precision and recall. The formula is defined as follows:

$$\begin{aligned} F1\text{–}score &= 2 * \frac{Precision * Recall}{Precision + Recall} \\ &= \frac{TP}{TP + \frac{1}{2}(FP + FN)} \end{aligned} \quad (5)$$

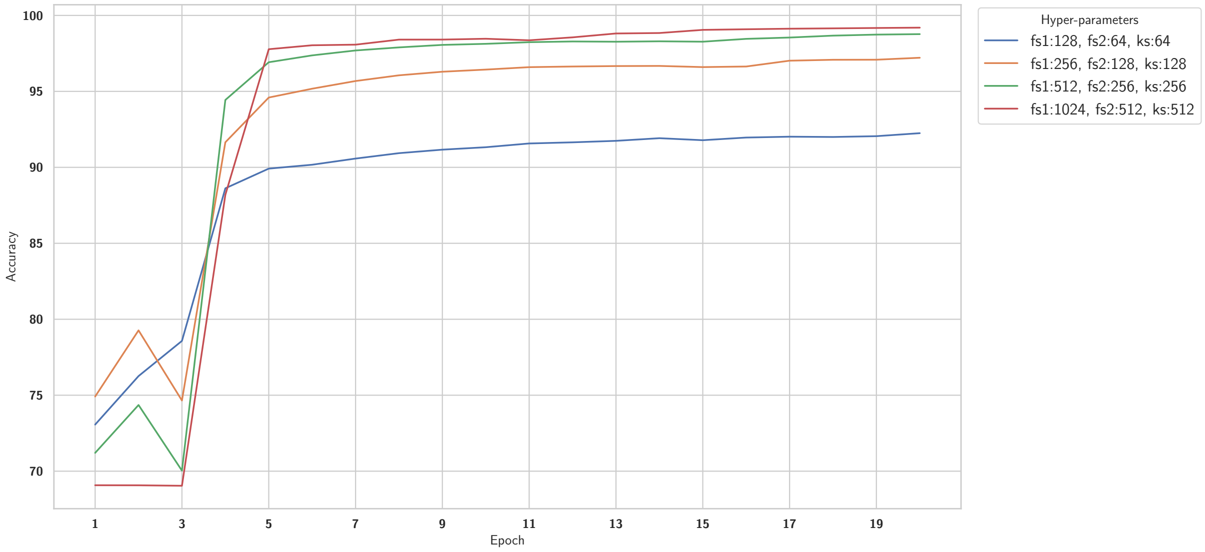


**Fig. 2** The effect of different hyper-parameters of DEEP*HLA on average accuracy validation sets in 3-fold cross validation on HLA dataset, where MissR is 50%. Since we tried to follow the exact choice of filter and kernel size from the original paper, the only main variable among these hyper-parameters is fs1 which is #Filters 1, representing the number of filters in the first convolution layer. The number of filters in the second layer (fs2) and kernel size (ks) are simply factors of the aforementioned hyper-parameter.

**Table 8** Average accuracy of different model on deletions in chromosome 22 of 1000 Genomes dataset over maximum linkage disequilibrium (LD) bins and MissR using 3-fold cross validation.

| | | Method | | | | | | |
|---|---|---|---|---|---|---|---|---|
| Missing Rate | LD | AE | SCDA+ | DEEP*HLA | Beagle | Minimac | STI-NE | STI |
| 0.05 | [0, 0.2) | 96.29(1.1e+00) | 96.31(1.2e+00) | 92.34(4.7e+00) | 95.16(1.5e-01) | 95.20(9.3e-02) | 97.22(1.1e-01) | 97.32(1.4e-01) |
| | [0.2, 0.4) | 98.78(4.0e-01) | 98.58(7.2e-01) | 95.07(5.8e+00) | 98.30(1.9e-01) | 98.36(1.4e-01) | 99.18(1.2e-01) | 99.32(3.4e-02) |
| | [0.4, 0.6) | 99.96(6.9e-02) | 99.84(1.4e-01) | 94.21(1.0e+01) | 99.96(6.9e-02) | 99.96(6.9e-02) | 99.96(6.9e-02) | 99.96(6.9e-02) |
| | [0.6, 0.8) | 99.76(1.0e-01) | 99.44(6.1e-01) | 93.63(1.0e+01) | 99.78(1.4e-01) | 99.90(6.9e-02) | 99.82(1.0e-01) | 99.88(2.1e-01) |
| | [0.8, 1] | 97.34(2.1e+00) | 98.24(9.2e-01) | 89.16(1.0e+01) | 98.46(3.5e-02) | 99.56(1.9e-01) | 99.02(2.5e-01) | 99.58(2.4e-01) |
| 0.1 | [0, 0.2) | 96.01(8.9e-01) | 96.63(3.7e-01) | 93.34(2.8e+00) | 95.02(7.4e-02) | 95.08(1.0e-01) | 96.85(5.7e-02) | 96.91(3.8e-02) |
| | [0.2, 0.4) | 98.81(4.5e-01) | 98.92(2.5e-01) | 96.29(3.7e+00) | 98.27(2.4e-01) | 98.49(1.6e-01) | 99.25(1.3e-01) | 99.30(7.5e-02) |
| | [0.4, 0.6) | 99.95(4.6e-02) | 99.93(2.3e-02) | 95.97(6.9e+00) | 99.95(2.3e-02) | 99.95(2.3e-02) | 99.95(6.1e-02) | 99.96(4.0e-02) |
| | [0.6, 0.8) | 99.60(1.1e-01) | 99.60(1.4e-01) | 95.90(6.4e+00) | 99.80(4.0e-02) | 99.91(6.1e-02) | 99.67(6.1e-02) | 99.83(9.2e-02) |
| | [0.8, 1] | 97.51(2.2e+00) | 98.46(3.5e-01) | 91.86(5.4e+00) | 97.90(6.0e-02) | 99.39(5.0e-02) | 98.68(1.0e-01) | 99.45(2.8e-01) |
| 0.2 | [0, 0.2) | 95.43(6.4e-01) | 96.37(7.5e-02) | 93.76(2.3e+00) | 95.17(6.4e-02) | 95.20(7.2e-02) | 96.44(5.5e-02) | 96.50(5.8e-02) |
| | [0.2, 0.4) | 98.55(2.9e-01) | 98.97(1.6e-02) | 96.86(2.8e+00) | 98.23(8.7e-02) | 98.38(8.6e-02) | 99.06(5.3e-02) | 99.08(3.6e-02) |
| | [0.4, 0.6) | 99.94(3.5e-02) | 99.94(3.5e-02) | 97.42(4.4e+00) | 99.93(2.3e-02) | 99.91(1.1e-02) | 99.94(4.0e-02) | 99.95(3.0e-02) |
| | [0.6, 0.8) | 99.70(7.2e-02) | 99.71(7.6e-02) | 97.15(4.3e+00) | 99.75(3.1e-02) | 99.85(2.3e-02) | 99.73(6.4e-02) | 99.86(6.9e-02) |
| | [0.8, 1] | 95.87(2.3e+00) | 98.21(1.8e-01) | 92.95(2.8e+00) | 97.82(8.9e-02) | 99.25(7.9e-02) | 98.53(1.0e-01) | 99.17(2.2e-01) |

where TP, FP, and FN are True Positive(s), False Positive(s), and False Negative(s), respectively. F1-score is a well defined metric for classification problems and there are three variants to it. Micro, Macro, and weighted. Here we used weighted f1-score. To calculate a weighted f1-score, first f1-score is calculated per class (alleles in our case) and then weighted average between the classes is computed. The weights are proportional to the support (instances) of the ground truth for the class.

Last but not the least, we use $R^2$ to compare the competing methods. There are various definitions of $R^2$, and they can use either hard-call genotypes or the dosage. Here since some methods did not provide the dosage for the output, we decided to use a variation of $R^2$ defined on hard-call genotypes in [22], which is defined as follows:

**Table 9** Average accuracy for sporadic missingness over 0.2 MissR and 3-fold cross validations for human chromosome 22 SVs. Best results are highlighted in bold. CNV events are entirely multi-allelic, and STI/STI-NE outperform other methods with a considerable performance gap while the performance for bi-allelic events is quite similar.

| | | **Method** | | | | | |
|---|---|---|---|---|---|---|---|
| Dataset | SV type | AE | SCDA+ | DEEP*HLA | Beagle | STI-NE | STI |
| Chr22(LD) | ALU | 92.3(1.1e-01) | 92.5(3.0e-01) | 92.5(2.9e-01) | 92.4(4.3e-01) | **92.6(2.2e-01)** | 92.5(2.5e-01) |
| | CNV | 93.5(6.8e-01) | 94.5(7.3e-01) | 93.5(2.7e-01) | 94.4(3.2e-01) | 95.1(3.2e-01) | **95.2(2.7e-01)** |
| | DEL | 96.0(1.7e-01) | 96.3(1.2e-01) | 96.1(6.1e-02) | 96.2(6.9e-02) | **96.4(1.9e-02)** | 96.4(8.4e-02) |
| | DEL_ALU | 44.7(3.4e+00) | 48.6(4.0e-01) | 43.0(1.9e+00) | 50.4(1.8e+00) | 49.3(1.5e+00) | **52.8(5.8e-01)** |
| | DUP | 99.9(2.2e-01) | 99.9(2.2e-01) | 99.9(2.2e-01) | 99.8(1.9e-01) | 99.9(2.2e-01) | 99.9(2.2e-01) |
| | INS | 99.3(5.4e-01) | 99.3(5.4e-01) | 99.3(5.4e-01) | 99.3(5.4e-01) | 99.3(5.4e-01) | 99.3(5.4e-01) |
| | INV | **99.3(2.5e-01)** | 99.3(3.5e-01) | 99.3(2.8e-01) | 99.3(2.8e-01) | 99.3(2.8e-01) | 99.3(2.8e-01) |
| | LINE1 | **93.6(1.8e-01)** | 93.4(4.7e-02) | 93.6(2.4e-01) | 93.5(3.7e-01) | 93.6(2.7e-01) | 93.6(2.4e-01) |
| | SVA | 92.5(2.1e-01) | 92.4(9.3e-02) | 92.2(1.3e-01) | 92.5(2.1e-01) | **92.6(1.1e-01)** | 92.3(4.5e-01) |
| Chr22(MAF) | ALU | 91.9(1.7e-01) | 92.0(1.1e-01) | 90.7(1.1e+00) | 91.7(1.8e-01) | 92.0(5.5e-02) | **92.1(1.9e-01)** |
| | CNV | 93.9(1.3e-01) | 95.0(1.5e-01) | 92.2(3.4e-01) | 94.3(3.9e-02) | 94.8(9.4e-02) | **95.1(4.7e-02)** |
| | DEL | 96.1(1.0e-01) | 96.4(3.6e-02) | 95.3(4.7e-01) | 96.1(3.7e-02) | 96.3(3.2e-02) | **96.4(1.0e-02)** |
| | DEL_ALU | 45.8(2.1e+00) | 51.4(6.3e-01) | 43.4(3.1e+00) | 49.0(1.5e-01) | 48.9(6.4e-01) | **52.8(4.1e-01)** |
| | DUP | 99.9(1.1e-01) | 99.8(2.0e-01) | 99.3(6.4e-01) | 99.9(1.1e-01) | 99.9(1.1e-01) | 99.8(4.0e-04) |
| | INS | 99.4(4.1e-01) | 99.4(4.1e-01) | 99.0(8.6e-01) | 99.3(3.9e-01) | 99.4(4.1e-01) | 99.4(4.1e-01) |
| | INV | **98.9(3.8e-01)** | 98.9(4.5e-01) | 98.4(1.1e+00) | 98.9(4.5e-01) | 98.9(4.5e-01) | 98.9(4.5e-01) |
| | LINE1 | 93.7(5.5e-01) | 93.8(9.4e-02) | 92.9(1.2e+00) | 93.5(3.9e-01) | 93.7(2.3e-01) | **94.0(2.6e-01)** |
| | SVA | 91.9(2.3e-01) | 91.9(2.1e-01) | 91.1(6.3e-01) | 91.8(3.0e-01) | 91.9(9.2e-02) | **92.0(2.2e-01)** |

**Table 10** Experimental results for imputing systematic missingness in chromosome 22 of human 1000 Genomes Project dataset. STI is trained using 0.8 MaskR. Metrics in this table use an identical categorical value to encode heterozygous alternative alleles. Bold values indicate top results in each row. In this dataset, Beagle outperforms other methods, and among DL methods RNN-IMP appears as close competitor to Beagle.

| Metric | Beagle | Minimac | RapidAE | RNN-IMP | STI |
|---|---|---|---|---|---|
| Accuracy | **97.2(1.1e+00)** | 96.6(1.3e+00) | 93.6(2.2e+00) | 97.1(1.1e+00) | 96.1(1.4e+00) |
| F1-score | **0.972(1.04e-02)** | 0.965(1.27e-02) | 0.932(2.23e-02) | 0.970(1.09e-02) | 0.959(1.32e-02) |
| IQS | **0.945(4.45e-03)** | 0.932(4.57e-03) | 0.863(1.88e-02) | 0.942(5.12e-03) | 0.917(1.67e-02) |
| $R^2$ | **0.938(1.12e-02)** | 0.920(1.19e-02) | 0.832(3.00e-02) | 0.935(1.20e-02) | 0.901(2.63e-02) |

$$R_i^2 = \frac{Cov^2(A_i, B_i)}{Var(A_i)Var(B_i)} \tag{6}$$

where $i$ is a single locus index and A and B are vectors of imputed and ground truth genotypes at a given locus $i$, respectively. The values in A and B are homozygous reference allele, heterozygous alternative allele, or homozygous alternative allele encoded as 0, 1, and 2 respectively.

**Acknowledgements.** This work is partially supported by the US National Science Foundation (DBI 1750632) and the National Institutes of Health (GM-0126567-03). This research includes calculations carried out on HPC resources supported in part by the National Science Foundation through major research instrumentation grant number 1625061 and by the US Army Research Laboratory under contract number W911NF-16-2-0189. We appreciate the suggestion provided by Dr. Francisco McGee in designing the model, leading to improvements in performance. Additionally, we would like to thank Dr. Kaname Kojima for helping us obtain the data for the Missing variant experiment and providing experimental results for RNN-IMP (the experiment in the supplement) and Emily Thyrum for proofreading the manuscript.

**Table 11** Inference time for the competing method in seconds. For the HLA dataset, STI is using a small batch size due to model size restrictions. With a slightly lighter model, the batch size could be doubled and the time would decrease to 9 seconds for the HLA set for STI.

| | Dataset | | | | |
|---|---|---|---|---|---|
| Method | Yeast | HLA | Chr-22 DEL | Chr-22 ALL | Omni2.5 |
| AE | 4 | 3 | $< 1$ | $< 1$ | N/A |
| SCDA+ | 4 | 3 | $< 1$ | $< 1$ | N/A |
| RapidAE | N/A | N/A | N/A | N/A | $< 1$ |
| DEEP*HLA | 33 | 4 | $< 1$ | $< 1$ | N/A |
| Minimac4 | N/A | 5 | 2 | N/A | 20 |
| Beagle5.4 | N/A | 36 | 8 | 12 | 4 |
| STI | 13 | 19* | 2 | 2 | 5 |

### Declarations

Not applicable.
